# An Epigenetic LTR-retrotransposon insertion in the upstream region of *BnSHP1.A9* controls quantitative pod shattering resistance in *Brassica napus*

**DOI:** 10.1101/858407

**Authors:** Jia Liu, Rijin Zhou, Wenxiang Wang, Hui Wang, Yu Qiu, Raman Rosy, Desheng Mei, Raman Harsh, Qiong Hu

## Abstract

Seed loss resulting from pod shattering is a major problem in oilseed rape (*Brassica napus* L.) production worldwide. However, the molecular mechanisms underlying pod shatter resistance are not well understood. Here we show that the pod shatter resistance at quantitative trait locus, qSRI.A9.1 is controlled by a *SHATTERPROOF1* (*SHP1*) paralog in *B. napus* (*BnSHP1.A9*). Expression analysis by quantitative RT-PCR showed that Bn*SHP1.A9* was specifically expressed in flower buds, flowers and developing siliques in the oilseed rape line (R1) carrying the qSRI.A9.1 allele with negative effect, but not expressed in any tissue of the line (R2) carrying the positive effect qSRI.A9.1 allele. Transgenic plants constitutively expressing Bn*SHP1.A9* alleles from pod resistant and pod shattering parental lines showed that both alleles are responsible for pod shattering via promoting lignification of en*b* layer, which indicated allelic difference of Bn*SHP1.A9* gene *per se* is not the causal factor of the QTL. The upstream sequence of Bn*SHP1.A9* in the promotor region harboring highly methylated long terminal repeat retrotransposon insertion (LTR, 4803bp) in R2 repressed the expression of Bn*SHP.A9,* and thus contributed to the positive effect on pod shatter resistance. Genetic and association analysis revealed that the *copia* LTR retrotransposon based marker *BnSHP1.A9*-_R2_ can be used for breeding for pod shatter resistant varieties and reducing the loss of seed yield in oilseed rape.

## Introduction

Oilseed rape (*Brassica napus* L.) is not only a major source of edible vegetable oil for human consumption, but also provides an important energy resource for stock-feed and biodiesel production. Upon maturity, siliques (pods) of oilseed rape open, dehiscing seed and causing significant yield loss; particularly if oilseed is harvested after the full maturity (BBCH scale 95). Pod shattering usually accounts for an approximately 10% yield loss on an average, however in certain environments it can cause yield loss up to 50% (Kadkol et al., 1984; Wang et al., 2007). In order to reduce such loss, oilseed rape is either harvested manually or mechanically before full maturation of seeds (windrowing). Premature harvest can reduce yield loss, but immature seed can lead to lower oil content (Østergaard et al., 2006) and higher chlorophyll content. In recent years, due to the shortage and high cost of labour, broad-acre cropping of oilseed rape has been preferentially combine harvested by farmers. Varieties with improved pod shatter resistance, amenable for mechanical harvesting provide a cost-effective solution for commercial oilseed rape production worldwide.

Genetic variation for pod shatter resistance exists in *Brassica napus*, *Brassica rapa*, *Brassica juncea* and *Brassica carinata* germplasm (Hu et al., 2012; Liu et al., 2016; Raman et al., 2014; Raman et al., 2017; Kadkol et al., 1984), and can be exploited to breed commercial varieties with improved pod shatter resistance. Genetic studies have revealed that pod shatter resistance is controlled by multiple genes. Several quantitative trait loci (QTL) associated with this trait have been localized on genetic and physical maps of oilseed rape (Hu et al., 2012; Raman et al., 2014; Liu et al., 2016). However, the identification of corresponding genes underlying QTL for pod shatter resistance in *B. napus* has not been reported yet.

In the model plant *Arabidopsis thaliana,* which belongs to the same Brassicacae family as oilseed rape, gene network involved in pod development and dehiscence has been elucidated (Liljegren et al., 2004). For example, MADS–box transcription factors *SHATTERPROOF1* (*SHP1*) and *SHATTERPROOF2* (*SHP2*) (Liljegren et al., 2000), basic helix-loop-helix (bHLH) gene *INDEHISCENT* (*IND*) (Liljegren et al., 2004), and *ALCATRAZ* (*ALC*) (Rajani and Sundaresan, 2001) regulate the differentiation of the dehiscence zone. The activity of valve margin identity genes is repressed by *FRUITFULL* (*FUL*) in the valves (Gu et al., 1998), and *REPLUMLESS* (*RPL*) in the replum (Roeder et al., 2003). Several phytohormones such as auxin, cytokinin and gibberellin are also essential for pod development and the expression of valve margin identity and *IND* genes (Larsson et al., 2014; Simonini et al., 2017; Arnaud et al., 2010; Zuniga-Mayo et al., 2014). Braatz et al (2018a and 2018b) have shown that induced mutations in *IND* and *ALC* homologues are linked with pod shatter resistance in oilseed rape. Ectopic expression of *FUL* gene from *Arabidopsis* has been shown to result in pod shatter resistance via inhibiting *SHP* expression in *B. juncea* (Ostergaard et al., 2006). However for commercial production of ‘conventional’ oilseed rape, a fine tuning of these genes is required to develop a desirable level of dehiscence.

Comparative mapping studies have shown that the *SHATTERPROOF* paralogs of *A. thaliana* (*SHP1* and *SHP2*) are located in the vicinity of the QTL associated with pod shatter resistance on chromosome A9 (designated as *BnSHP1.A9* and *BnSHP2.A9*) in Australian and Chinese oilseed rape populations, including the R1/R2 DH population utilized in this study (Liu et al., 2016; Raman et al., 2014). *SHP1* and *SHP2* are MADS-box genes, which regulate cell differentiation of the valve margin and promote lignification (Liljegren et al., 2000). Both genes are highly homologous and functionally redundant.

In the present study, we cloned the *BnSHP1.A9* gene underlying the resistant QTL qSRI.A9.1 for pod shatter resistance and characterized its functional role by employing comparative expression analysis using quantitative RT-PCR, anatomical and transgenic approaches. Our findings suggested that the LTR retroelement insertion silences the expression of *BnSHP1.A9* epigenetically via DNA methylation, and contributes to the pod shatter resistance in oilseed rape.

## Materials and Methods

### Plant materials and evaluation of resistance to pod shattering

For genetic analysis we used parental lines, R1 (pod shatter resistant), R2 (pod shatter prone) and a mapping population comprising of 96 DH lines that showed segregation for pod shatter resistance, developed from a cross between R1 and R2 (Liu et al., 2013). In addition, we used a total of 135 diverse accessions, comprised of four winter type, 119 semi-winter type and 12 spring type (Supplemental table 1) to validate the association between pod shatter resistance index (PSRI) and the *BnSHP1.A9* promoter specific marker. All accessions were evaluated for pod shatter resistance using random impact test as described previously (Liu et al., 2016).

### Sequence analysis of *BnSHP1.A9*

Total DNA was extracted from the leaves of 4 weeks old seedlings by CTAB method (Stein et al., 2001). The reference sequence of *BnSHP1.A9* (BnaA09g55330D) from the *B. napus cv.* Darmor genome (www.genoscope.cns.fr/brassicanapus) was used as a template to design specific primer pairs (Table1) for cloning the genomic sequence and open reading frame of *BnSHP1.A9*. DNA amplification was performed in a 20 μl reaction system comprising 2 μl (10 μM) of forward and reverse primers, 4 μl 5× TransStart^®^FastPfu Fly buffer, 2 μl 2.5mM dNTPs, 2 l 5× PCR stimulant, 0.4 μl 50mM MgSO_4_, 1 U TransStart FastPfu Fly DNA Polymerase (Beijing TransGen Biotech Co., Ltd., Product Code: AP231), and 20-30 ng template DNA. After initial denaturation of template DNA at 95 °C for 3 min, PCR was carried out following 34 cycles of 20 seconds at 94 °C, 30 seconds at about 5 °C lower than melting temperature of the primer pair used, 1 min/kb at 72 °C with a final extension of 5 min at 72 °C. The PCR products were separated by electrophoresis in 0.8% low melting agarose gels, the target bands were purified, cloned into the pEASY-Blunt Zero cloning vector (Beijing TransGen Biotech Co. Ltd., Product Code: CB501-02) and then transformed into *Trans*1-T1 competent *E. coli* cells by the heat shock method (Beijing TransGen Biotech Co. Ltd., Product Code: CB501, CT101). Four to six positive clones were randomly selected and sequenced with M13 sequencing primers at Shanghai Sangon Biotech Co. Ltd (http://www.sangon.com/). Sequences were then analyzed by MultAlin online software (Corpet, 1988). To determine the physical location and sequence similarities of *BnSHP1.A9* clones, the BLASTn was used against the Darmor reference sequence (http://www.genoscope.cns.fr/brassicanapus/). To clone the upstream sequence of *BnSHP1.A9,* we developed two specific primer pairs for R1 (BnSHP1.A9p1) and R2 (BnSHP1.A9p2) that are listed in Table 2.

**Table 1.**
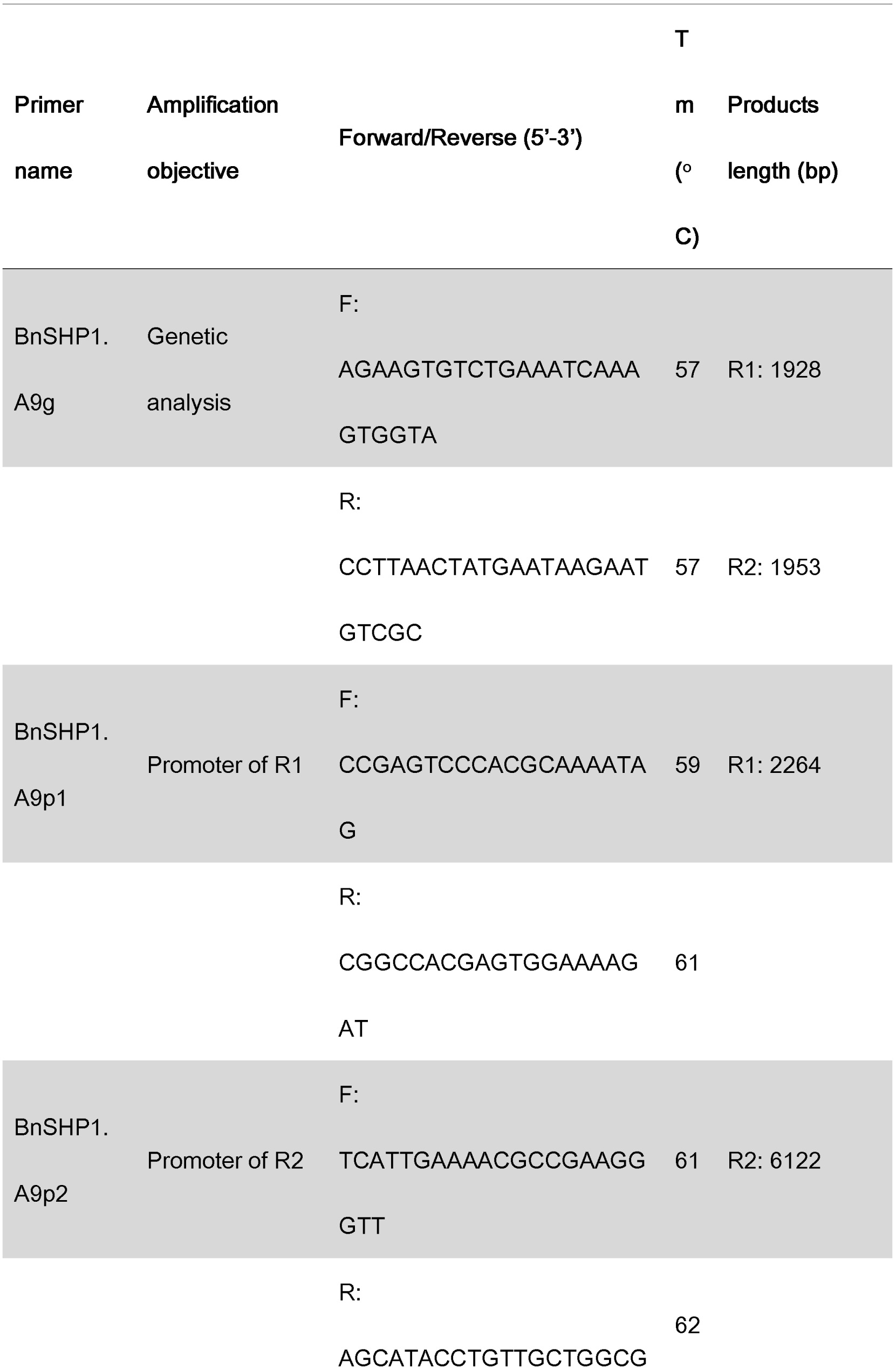
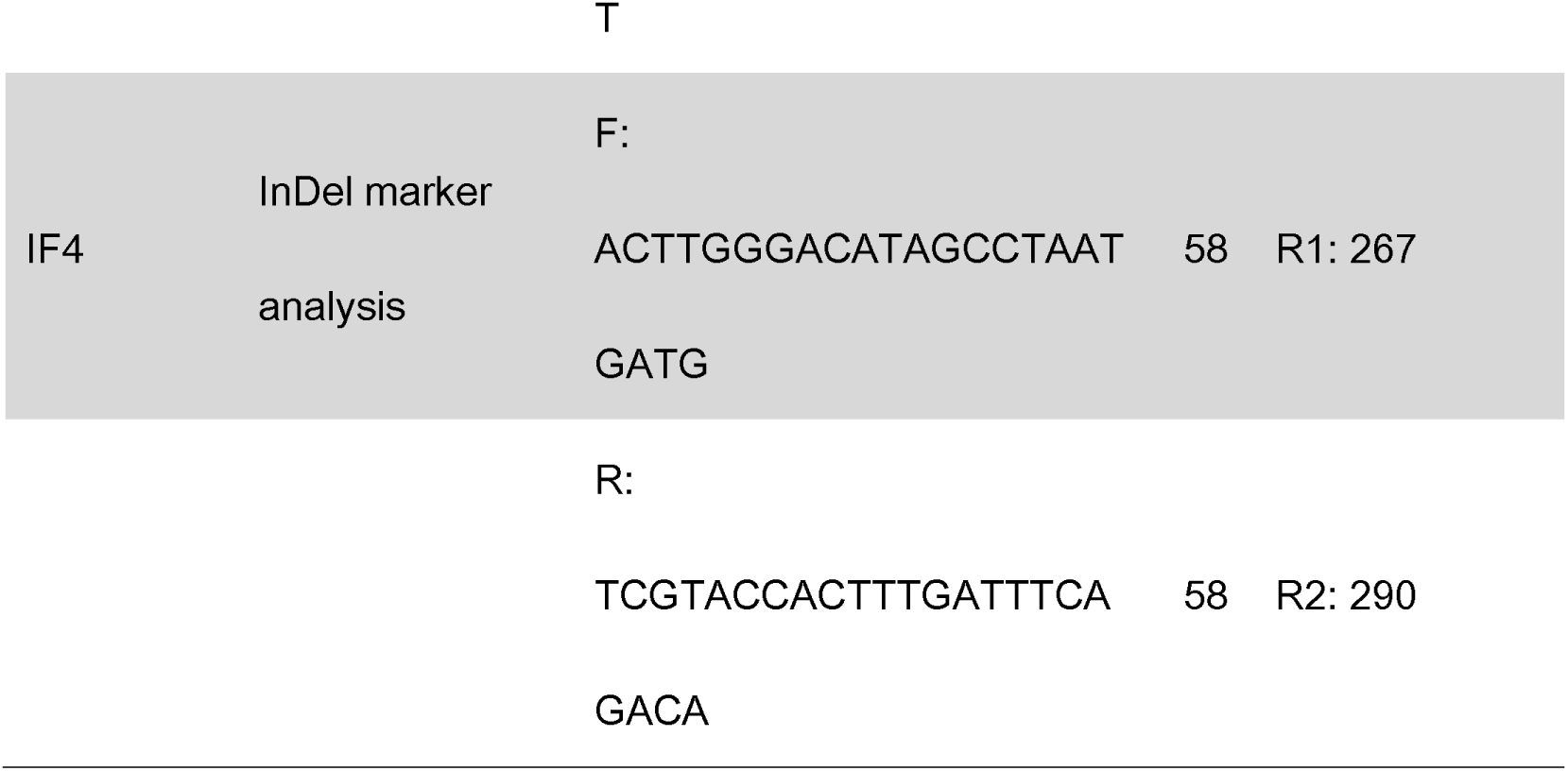
Primer pairs used for *BnSHP1.A9* gene cloning and genetic analyses

**Table 2.**
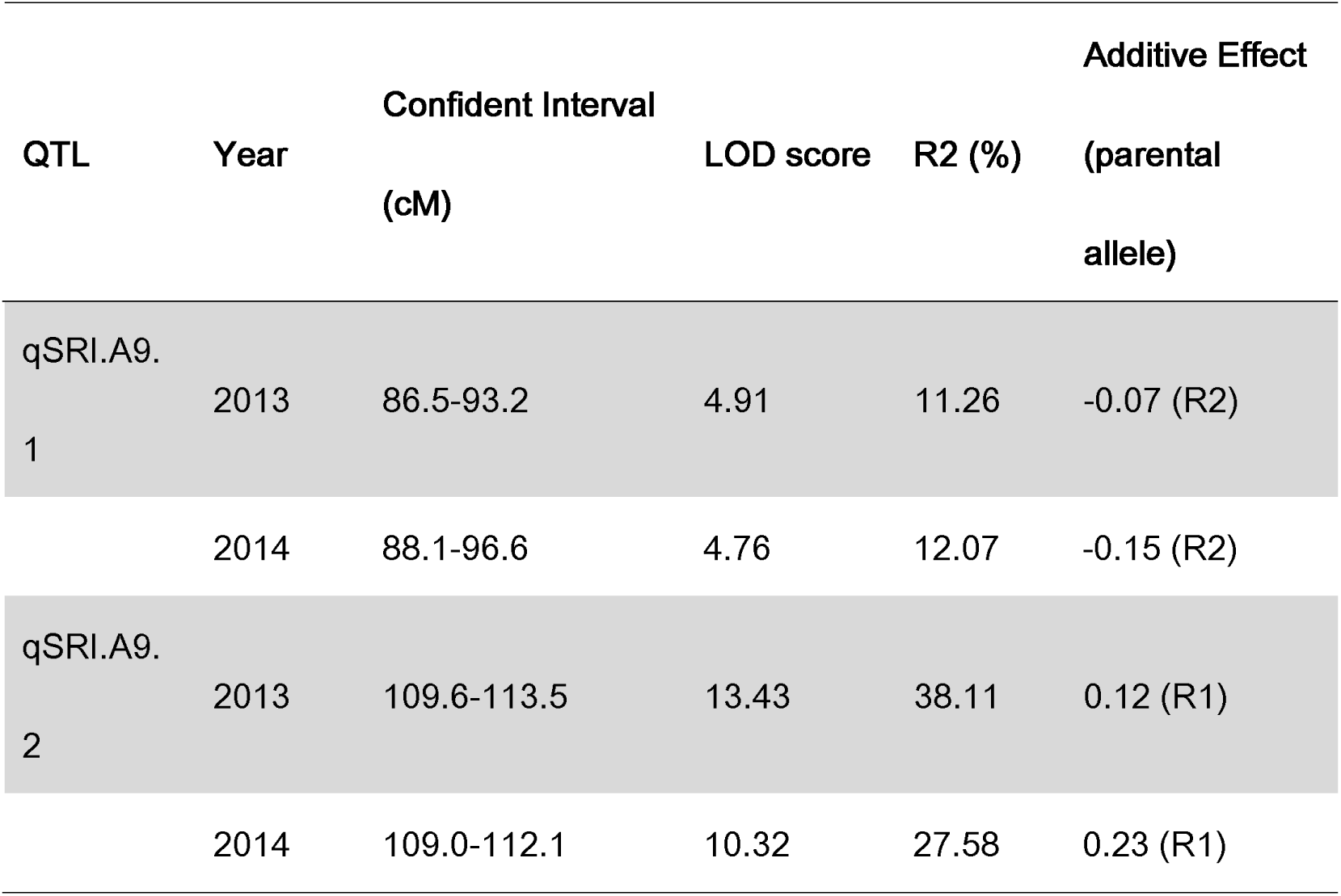
QTL associated with pod shatter resistance in a doubled haploid population from R1 (pod shatter resistant)/R2 (pod shatter susceptible). DH lines were evaluated for resistance using random impact test. R^2^ refers to phenotypic variance explained

| QTL | Year | Confident Interval<br>(cM) | LOD score | R2 (%) | Additive Effect<br>(parental<br>allele) |
| --- | --- | --- | --- | --- | --- |
| qSRI.A9.<br>1 | 2013 | 86.5-93.2 | 4.91 | 11.26 | -0.07 (R2) |
|  | 2014 | 88.1-96.6 | 4.76 | 12.07 | -0.15 (R2) |
| qSRI.A9.<br>2 | 2013 | 109.6-113.5 | 13.43 | 38.11 | 0.12 (R1) |
|  | 2014 | 109.0-112.1 | 10.32 | 27.58 | 0.23 (R1) |

### Development and assay of *BnSHP1.A9*-specific marker

The genomic sequences of *BnSHP1.A9* from R1 and R2 were aligned in MEGA7 (Kumar et al., 2016) using both the reference genomic and coding sequences (BnaA09g55330D) of Darmor-*bzh*. Based on the sequence differences we developed a co-dominant Indel marker (*BnSHP1.A9*-IF4) targeting the indel difference of the first intron of *BnSHP1.A9* between the parental lines of the R1/R2 population for genetic mapping and the allelic diversity analysis (Table 1). PCR was performed in a volume of 10 μl system, including 5 μl 2×Taq MasterMix, 1 μl (10 μM) of each primer, 2 μl ddH_2_O, 1 μl genomic DNA (20-30 ng). The PCR condition used was as following: 3 min at 94 °C, 35 cycles of 30 s at 94 °C, 30 s at 57 °C, 30 s at 72 °C, with a final extension of 5 min at 72 °C. The PCR products were examined on either 3% agarose gel by electrophoresis at 130 V for 1 hour or using capillary electrophoresis on an automated CEQ2000 system (Beckman-Coulter) as described previously (Raman et al., 2005). The gels were stained with Syber-green and visualized under UV light. The PCR yielded a 267-bp product from R1 (pod shatter resistance allele) and a 290-bp product from R2 (pod shatter prone allele).

### QTL mapping

The 96 DH lines derived from R1/R2 cross were genotyped with an InDel marker *BnSHP1.A9-IF4* and other new InDel and SSR makers developed based on sequence variation between the two parental lines in the vicinity of qSRI.A9 region (Supplemental table 2). This polymorphic data was integrated with the Illumina Brassica 60K Infinium ® SNP array and pod shatter resistance data generated in our previous study (Liu et al., 2016). To determine putative QTL for pod shatter resistance, we performed the composite interval mapping (CIM) using WinQTL Cartographer 2.50 (Zeng, 1994; Wang et al., 2006). The genome scan was performed at every 2 cM to estimate the likelihood of a QTL and its corresponding phenotypic effect (R^2^). The empirical threshold was computed using 1,000 permutations (overall error level 5%) as described in Churchill and Doerge (1994).

### Analysis of *BnSHP1.A9* transcript levels

To detect the dynamic expression patterns of the *BnSHP1.A9* gene, tissues of root, stem and young leaf were collected from 4 weeks old plants, while bud, fully-open flower, and developing pods were collected at 10, 20, 30 and 40 days after flowering (DAF) from R1 and R2 parental and DH lines (DH56 and DH82), and snap frozen in liquid nitrogen. For each sample, including pods at different developmental stages, three biological replicates were used for semi-quantitative reverse transcription PCR (RT-PCR). Total RNA was extracted using the RNAprep Pure Plant Kit (TIANGEN Biotech Beijing Co., Ltd. Product code: DP432). DNAse I-treated RNA was reverse transcribed using the cDNA synthesizing kit following the manufacturer’s instructions (TIANGEN Biotech Beijing Co., Ltd. Product code: KR106-02). To eliminate interference from other paralogues, an allele-specific primer pair *BnSHP1.A9-*RT (F/R) was designed (Supplemental table 3) and the specificity was validated by PCR product sequencing. The a*ctin* gene was used as an internal control for semi-quantification of relative expression values.

### DNA methylation analysis

McrBC endonuclease (NEB, M0272S) was firstly employed to analyze the DNA methylation status of *BnSHP1.A9* promoter region and genic region. 0.5 µg genomic DNA extracted from 20 DAF siliques of R1 and R2 was digested respectively with 5 U McrBC overnight at 37 °C, the methylated plasmid DNA in the product was used as positive control. The McrBC-digested DNA was then used as template to amplify *BnSHP1.A9* promoter and genic region with allele specific primer pairs (Supplemental table 4). The Chop PCR products were then checked by agarose gel electrophoresis and stained with Syber-green and visualized under UV light.

Detailed analysis of promoter DNA methylation was then performed with bisulfite sequencing method (Gruntman et al., 2008). Genomic DNA of R1 and R2 from 20 DAF siliques was bisulfite treated using EpiTect Bisulfite Kit (Qiagen 59104) following the instructions of manufacturer. Bisulfite sequencing primers were designed with MethPrimer online tool (http://www.urogene.org/cgi-bin/methprimer/methprimer.cgi) and primer designing tool in NCBI. The bisulfate-treated DNA was used as template to amplify the target fragments of R1/R2 promoter with specific bisulfite primers (Supplemental table 5) and the resulting PCR fragments were cloned into pTOPO-T simple vector (Aidlab CV1501). At least 24 positive clones were sequenced from each fragment. Vector sequences together with primer sequences of sequencing results were first trimmed out and the remaining sequences were then blasted against *Brassica napus* reference genome database (http://www.genoscope.cns.fr/brassicanapus/) to confirm the specificity. The methylation status of the specifically amplified fragments was then analyzed by Kismeth online tool (http://katahdin.mssm.edu/kismeth/revpage.pl). The data of methylated cytosines (CG, CHG and CHH) between the two parental lines were collected and compared by t-test.

### Generation and analysis of transgenic lines overexpressing *BnSHP1.A9*

To examine the function of different *BnSHP1.A9* alleles from R1 and R2 parental lines, two overexpression vectors driven by CAMV35S promoter, 35S::*BnSHP1.A9*-_R1_ and 35S::*BnSHP1.A9*-_R2_, were constructed by cloning *BnSHP1.A9* coding sequence from R1 and R2 into the pCAMBIA-1301 vector using *Nco* I-*BstE* II cloning sites. *BnSHP1.A9* cDNA was amplified from developing pods using primer pair BnSHP1.A9orf (Table 2). The two overexpression vectors were then transformed into *Agrobacterium tumefaciens* GV3101 strain individually and then used for transformation of the *B. napus* line R1 (resistant to pod shatter) by *A. tumefaciens* mediated method (Hood et al., 1993). Transgenic T0 and T1 plants were confirmed by PCR detection of hygromycin resistance gene with primer pair *HptII*-F/R (Supplemental table 6). The expression level of *BnSHP1.A9* in transgenic plants was evaluated by RT-PCR. Since CAMV35S promoter could drive constitutive expression of downstream target genes in all tissues of transgenic plant, for the convenience of RNA extraction and early detection, we used leaf tissue for determining the expression level of *BnSHP1.A9* in transgenic plant. Transgenic lines of the T1 generation derived from both constructs were evaluated for pod shatter resistance using the random impact test as described previously (Liu et al., 2016).

## Results

### Sequence variation of *BnSHP1.A9* underlying pod shatter resistance

We isolated the full length of *BnSHP1.A9* gene from both the R1 and R2 parental lines of the DH mapping population. The size of *BnSHP1.A9* genomic sequences varied from 2,737 bp in R1 to 2,757 bp in R2, with 7 exons and 6 introns (Figure 1C). Sequence alignments revealed polymorphisms in the form of SNPs and InDel between the two parental lines. There were 9 SNPs in the exon sequences, including 2 nonsynonymous SNPs in exon1 at #25 (T/G) and #203 (G/T), leading to tyrosine (Y) to aspartic acid (D) and arginine (R) to leucine (L) amino acid variation, respectively. The point mutation in exon 3 (#1751,T/C) causes Y to H (histidine) amino acid alteration, while A/T mutation at #2480 in exon 6 leads to D to valine (V) amino acid change (Supplemental table 7). However, the most abundant sequence divergence was found in the first intron including 21 SNPs and 10 InDels between R1 and R2, which enabled us to develop a gene specific marker for *BnSHP1.A9*.

**Figure 1.**
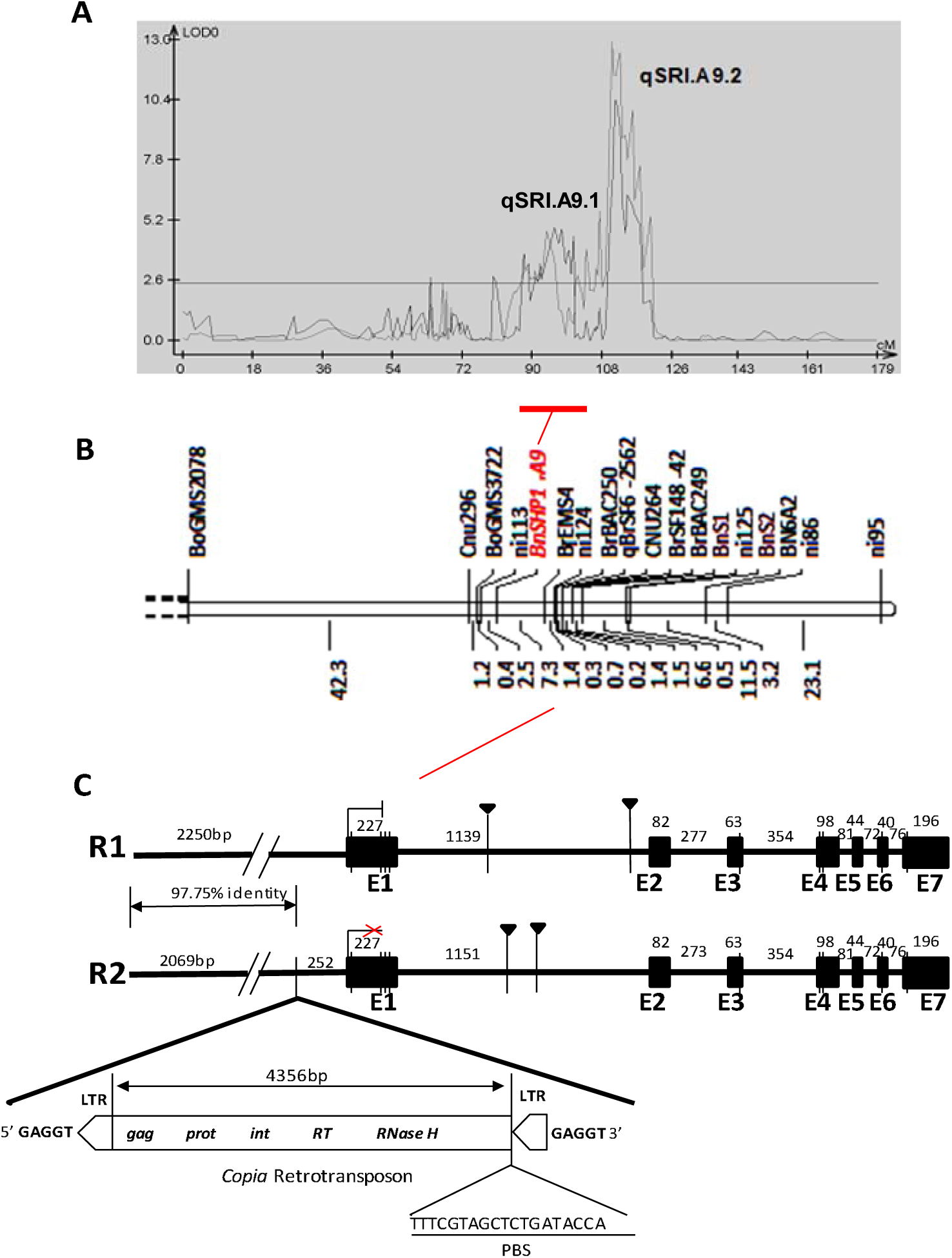
**Genetic mapping of the QTL; qSRI.A9.1 and underlying the candidate *SHATTERPROOF1*paralog of *A. thaliana* in *B. napus*, BnSHP1.A09. A. The schematic diagram of the LOD curve for the QTL of pod SRI of the A9 linkage group.** The abscissa coordinates represent to the genetic linkage group, and the ordinate represents to the LOD value. **B. Partial linkage map of chromosome A09, illustrating of the genetic position (cM) of *BnSHP1.A9* locus underlying qSRI.A9.1 for pod shatter in R1/R2 mapping population.** The left indicates the tag name (the newly developed Indel marker and SSR markers) and corresponding genetic map on the linkage group, corresponding to the location of the corresponding QTL for the pod SRI. **C. The structural variations present in the *BnSHP1.A9* gene between the parental lines of the DH mapping population R1 and R2. QTL was mapped using gene specific InDEL markers.**

In order to determine whether the polymorphism in *BnSHP1.A9* is associated with pod shatter resistance in a DH population derived from R1 and R2, we developed a primer pair (IF4) for specific amplification of *BnSHP1.A9* alleles (Table 1). IF4 amplified a 267-bp fragment from R1 and a 290-bp fragment from R2. Genotypic analysis of all the 96 DH lines showed a segregation for a single locus, with 35 lines containing the R1 allele and 55 lines containing the R2 allele (χ2 = 4.44, *P* = 0.035) (Supplemental table 8). These marker data were integrated with previously obtained SNP data from the R1/R2 DH population (Liu et al., 2016). QTL analysis identified two genomic regions, *qSRI.A9.1* and *qSRI.A9.2* for pod shattering resistance on chromosome A9, explaining 11.26% - 12.07% and 27.58% - 38.11% phenotypic variation, respectively (Figure 1A; Table 2). At *qSRI.A9.1*, the R2 allele contributed a positive effect to pod shatter resistance, whereas the R1 allele contributed a negative effect. In contrast, at *qSRI.A9.2*, the R1 allele contributed a positive effect to pod shatter resistance, whereas the R2 allele contributed a negative effect. Linkage analysis revealed the order of markers to be ni113 – *BnSHP1.A9* – BrEMS4; the *BnSHP1.A9* gene was mapped within the *qSRI.A9.1* genomic region for pod shatter resistance (Table 2, Figure 1A, B).

### A LTR retrotransposon is inserted in *BnSHP1.A9* promoter of R2

To gain an understanding on promoter region of the two *BnSHP1.A9* alleles, we sequenced and analyzed the upstream region of *BnSHP1.A9* from both parental lines of the mapping population. We identified a 4,803 bp long terminal repeat (LTR)/copia retrotransposon insertion at 252 bp upstream the start codon of *BnSHP1.A9* in R2 in opposite orientation (Figure 1C). This LTR retrotransposon contains the typical retro-transposable element structure, including a predicted 4,356 bp single open reading frame with conserved *gag*, *prot*, *int*, *RT* and *RNaseH* domains, two identical 169 bp 5’ and 3’ LTRs flanked by 5 bp direct repeat sequence (5’-GAGGT-3’) (Cao et al., 2015). Sequence alignment of this LTR retrotransposon against the NCBI database revealed 100% sequence identity with a *B. rapa* A9 scafford (LR031568.1), which suggested this LTR retrotransposon insertion may have been originated from *B. rapa* in R2 paternal line of R1R2 DH mapping population.

### Overexpression of *BnSHP1.A9* alleles promoted pod shattering in pod shatter resistant ‘R1’ line

Sequence analysis of *BnSHP1.A9* revealed 4 nonsynonymous SNPs in the coding sequence of *BnSHP1.A9* alleles of the two parental lines and three of them are located in the predicted conserved MADS-box and K-box domains. To verify whether these SNPs have any ‘phenotypic’ effect on gene function, we overexpressed *BnSHP1.A9* coding sequence (CDS) under the control of constitutive promoter from either R1 or R2 in pod shatter resistant R1 background respectively (named *BnSHP1.A9*-_R1_ *and BnSHP1.A9*_-R2_ hereafter). A total of 71 transgenic plants were generated and subsequently assessed for pod shatter resistance index by RIT. The PSRI of the transgenic T1 plants varied from 0.35-0.53, in comparison with the ‘wild-type (untransformed)’ R1 plants, which averaged 0.83 (n = 8). However, no statistically significant difference was found for PSRI between the transgenic *BnSHP1.A9*-_R1_ *and BnSHP1.A9*_-R2_ overexpression lines (OE). For example, four OE lines T18, T24, T26 and T33 of *BnSHP1.A9*-_R1_ had similar PSRI as that of the T9 line of *BnSHP1.A9*-_R2_ (Figure 2).

**Figure 2.**
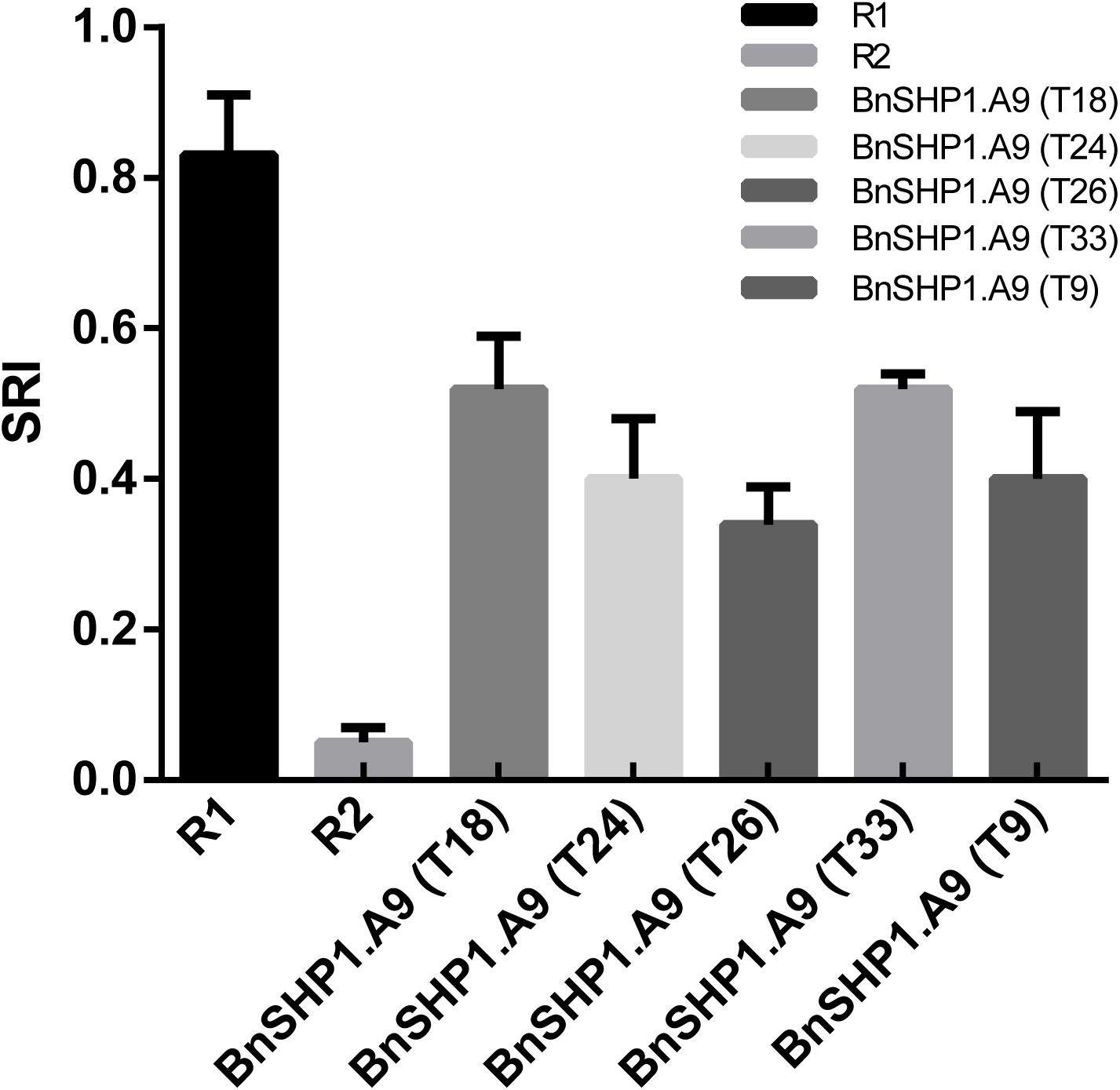
Pod shatter resistance index (PSRI) of different transgenic plants (T1) and two parental lines. PSRI was assessed with random impact test and 20 pods/each line were assessed with two replications. Standard deviation of each line is also shown.

Expression of *BnSHP1.A9* in developing pods in 15 selected transgenic T1 plants (three from each T1 line) were examined using semi-quantitative RT-PCR. All the tested T1 plants exhibited elevated level of *BnSHP1.A9* expression (Supplemental figure 1). These results proved that both alleles of *BnSHP1.A9* gene (R1 and R2) are functional in promoting pod shattering in *B. napus.* To explore whether overexpression of *BnSHP1.A9* causes dehiscence zone differentiation in siliques, we analyzed pod anatomical structure of *BnSHP1.A9*-_R1_ *and BnSHP1.A9*_-R2_ OE lines as described in Raman et al. (2017). A more compact arrangement of the ‘*en*b’ layer at the valve margin was observed in pod cross sections of the *BnSHP1.A9*-_R1_ and *BnSHP1.A9*_-R2_ OE T1 lines than in the wild type (Figure 3). Our results indicate that *BnSHP1.A9* gene promotes pod shattering through the increase of the number of lignified cells in the ‘*en*b’ layer, thus enhancing tension caused by differential contraction of the fruit wall tissue in oilseed rape.

**Figure 3.**
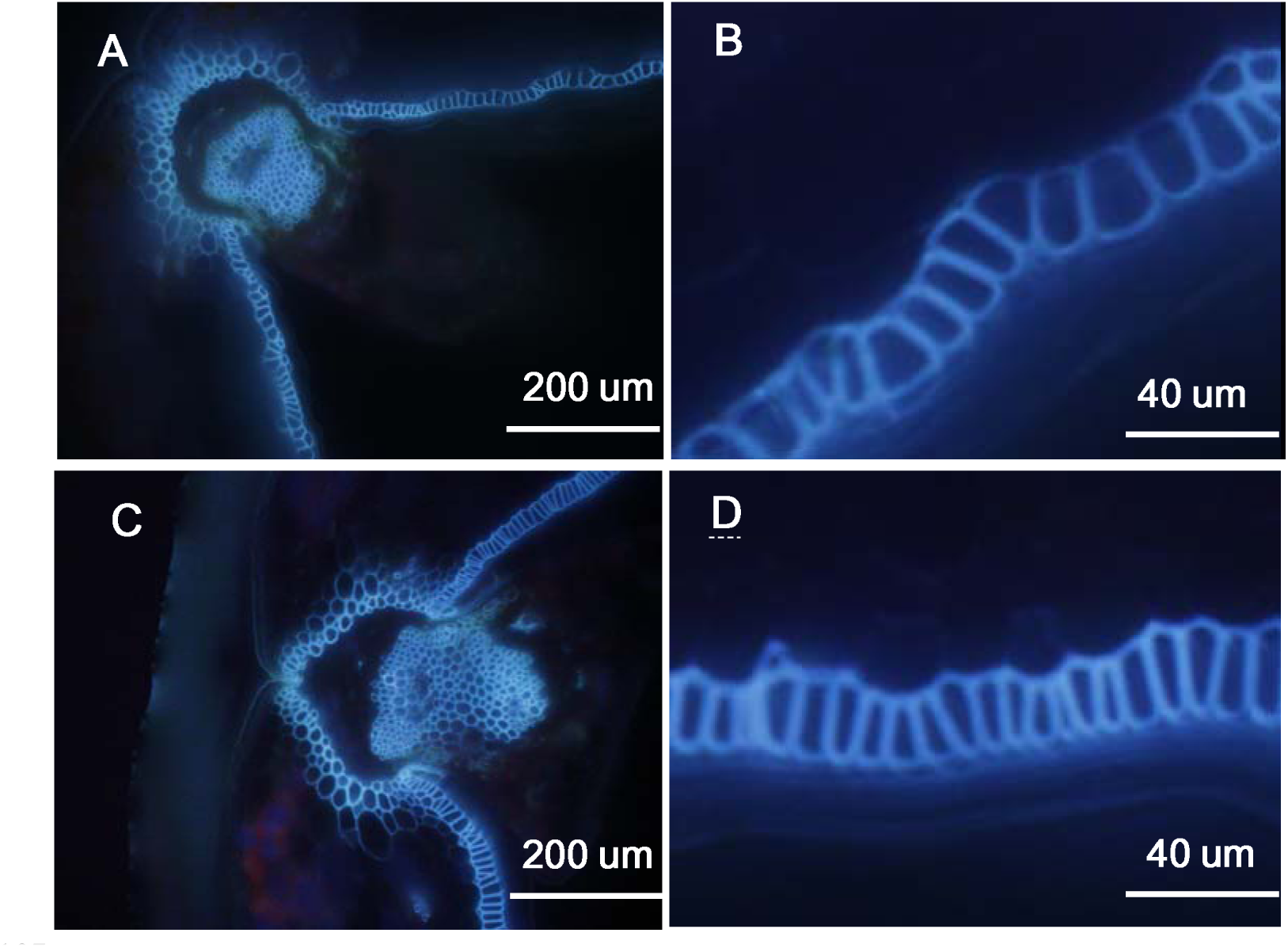
Histological analysis of 30-DAP silique cross-sections of 35S:*BnSHP1*.A9 transgenic rapeseed oil lines. (**A** and **B**) Microscopy observations of cross-sections of R1 siliques. Bars, 200 μm and 40 μm. (**C** and **D**) Microscopy observations of cross-sections of 35S:BnSHP1.A9 transgenic rapeseed oil siliques. Scale bars, 200 μm and 40 μm.

### Expression of *BnSHP1.A9* is repressed in R2

To test the dynamic expression of *BnSHP1.A9*, we investigated its expression pattern in root, stem, leaf, bud, flower, and developing siliques in R1 and R2 parental lines of the mapping population. Since there is a highly homologous copy of *SHP1* on chromosome C08 (BnaC08g29520D), to eliminate potential nonspecific amplification, we designed a specific RT-PCR primer pair for *BnSHP1.A9* based on SNPs between coding sequences of these two homologues. The result showed that *BnSHP1.A9* was expressed exclusively in bud, flower and developing siliques in R1. In contrast, almost no expression in either vegetative organs or reproductive organs could be detected in R2 (Figure 4A).

**Figure 4.**
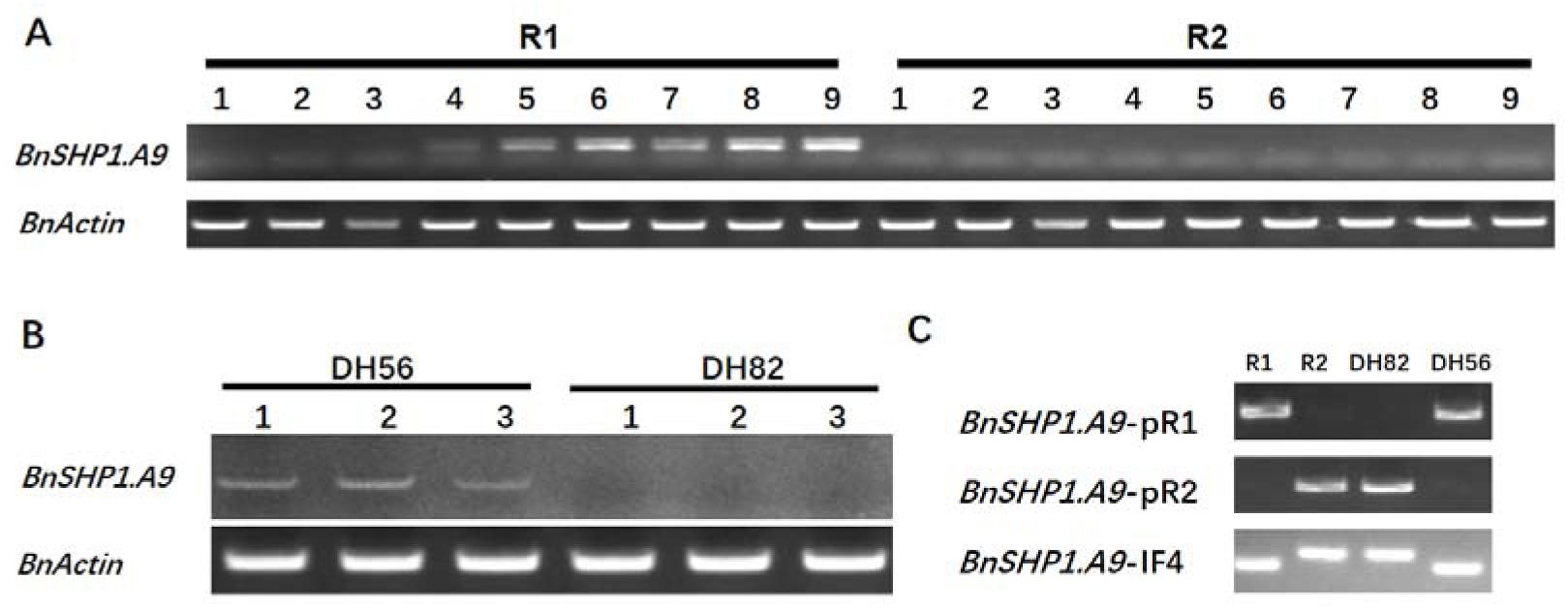
Expression status of *BnSHP1.A9* correlates with LTR retrotransposon insertion in parents and DH lines. **A**, *BnSHP1.A9* expression analysis of different tissues and silique, taken at different developmental stages of R1 and R2. *BnActin* gene was used as an internal control for relative expression analysis. 1: Root; 2: Stem; 3: Leaf; 4: Bud; 5: Flower; 6: 10d silique; 7: 20d silique; 8: 30d silique; 9: 40d silique. **B**, *BnSHP1.A9* expression status in DH56 and DH82 lines. 1: 10d silique; 2: 20d silique; 3: 20d silique. **C**: Genotyping of R1, R2, DH56 and DH82 with specific promoter primers of proR1, proR2 and InDel marker of *BnSHP1.A9*.

### The LTR insertion correlates with *BnSHP1.A9* repression in DH lines

Since the LTR element insertion locates in the upstream of the start codon of the R2 *BnSHP1.A9* allele, it may play a role in the repression of *BnSHP1.A9* expression. To further test this hypothesis, we investigated the 96 R1/R2 DH lines using promoter specific primers of *BnSHP1.A9* and selected two DH lines (DH56 and DH82 lines) that had the same genotype of *BnSHP1.A9* allele as R1 and R2, respectively. We then analyzed *BnSHP1.A9* expression level in 10, 20, 30 DAF siliques of DH56 and DH82 lines by RT-PCR. A weak band was visible in the siliques of all three developing stages of DH56, which does not contain the LTR insertion (LTR^-^). There was no amplification in any of the three silique samples in DH82 containing the LTR insertion (LTR^+^) (Figure 4B). On the basis of these data together with the expression pattern of *BnSHP1.A9* in the two parental lines, our results suggest that the LTR insertion in upstream of *BnSHP1.A9* is responsible for repression of *BnSHP1.A9* transcript.

### DNA methylation of LTR retroelement insertion is responsible for the repression of *BnSHP1.A9*

To understand the basis of LTR insertion mediated repression of *BnSHP1.A9*, we performed site-specific chop-PCR and bisulfite sequencing of genomic DNA extracted from 20 DAF silique of R1 and R2 to analyze DNA methylation status of the upstream and genomic regions of *BnSHP1.A9*. Chop-PCR amplified no bands from the LTR insertion fragments (b3, b4, b5 and b6) of R2 (Figure 5C and 5E) because of hypermethylation of the sequences, whereas same bands were amplified from other fragments of R2 as those from R1 (Figure 5B and 5D). These observations hinted that DNA methylation mainly occurred in the LTR insertion. The cytosine residues of LTR inner region were also decayed along far away the central insertion region.

The bisulfite sequencing results showed that the LTR retrotransposon and the transcription start region of *BnSHP1.A9* in R2 was hypermethylated, while the promoter region of *BnSHP1.A9* in R1 was hypomethylated (Figure 6B). The methylation level of mF4 region in R2 located at about 1.8 kb upstream the LTR element and the corresponding region of mF3 in R1 are much lower than the downstream regions close to the start codon (Fig. 6B and 6C). In mF1 region, we found the 100% cytosine (C) residues in all sites of CG and CHH (H rerpresents any residues other than G) were methylated in R2 because of the LTR insertion, whereas only 7.84% cytosine residues of the corresponding sites were methylated in R1. The mF2, mF3 and mF4 regions in R1 were hypomethylated from 2.37% to 6.10%. We investigated the methylation level of mF2 and mF3 of the LTR region in R2. In the mF2 region, 97.49% of cytosine residues was hypermethylated, while only 50.47% of cytosine residues were methylated in mF3 region. Among these cytosine residue containing sites in mF3 region, cytosine residues of CG were still hypermethylated, however the methylation of cytosine residues of CHG and CHH were significantly decreased to 43.85% and 17.53%. For mF4, the upstream region of about 2 kb away from the LTR and 7 Kb away from the transcription start codon of *BnSHP1.A9*, the methylation level was significantly decreased to a normal level of about 6.10% in both lines. Thus, methylation by the LTR retrotransposon in the vicinity of *BnSHP1.A9* promoter region in R2 may directly affect in the down-regulation of *BnSHP1.A9* expression.

**Figure 5.**
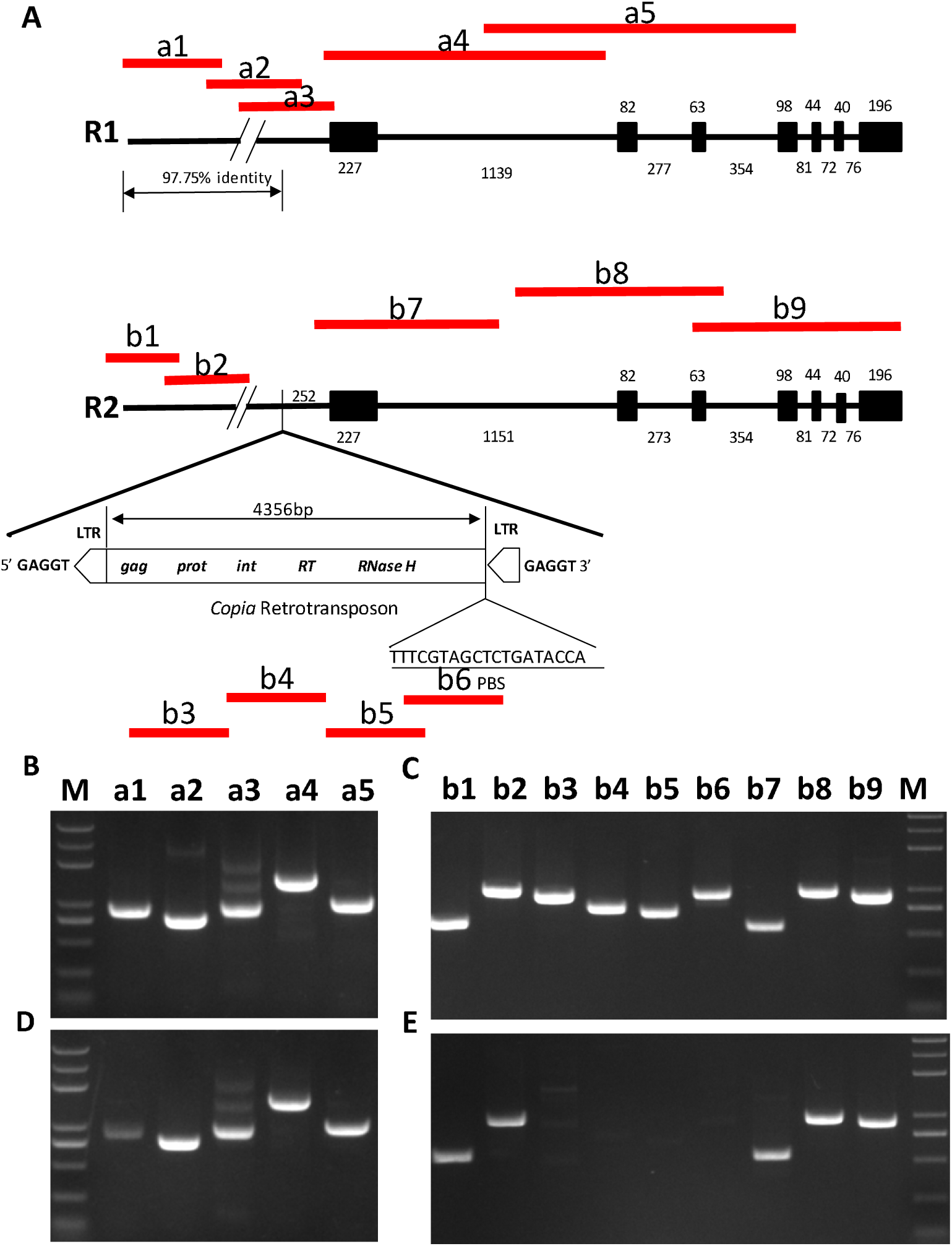
Cross validation of DNA methylation status of *BnSHP1.A9* in R1/R2 silique genomic DNA. **A,** amplified fragments illustrated in *BnSHP1.A9* DNA from R1/R2 **B, D**: chop-PCR with *BnSHP1.A9* gDNA template and McrBC digested gDNA of 20 DAF silique from R1 without McrBC digestion; a1: −2412/−1542, a2: −1616/−882, a3: −902/−30, a4: −49/+1413, a5: +1034/+2007 **C, E**: chop-PCR with *BnSHP1.A9* gDNA template and McrBC digested gDNA of 20 DAF silique from R2. b1: −7412/−6823, b2: −6753/−5765, b3: −3191/−2285, b4: −2538/−1788, b5: −1707/−1027, b6: −1035/−102, b7: −49/+436, b8: +1035/+2012, b9: +1883/+2793 M: Trans 2k plus.

**Figure 6.**
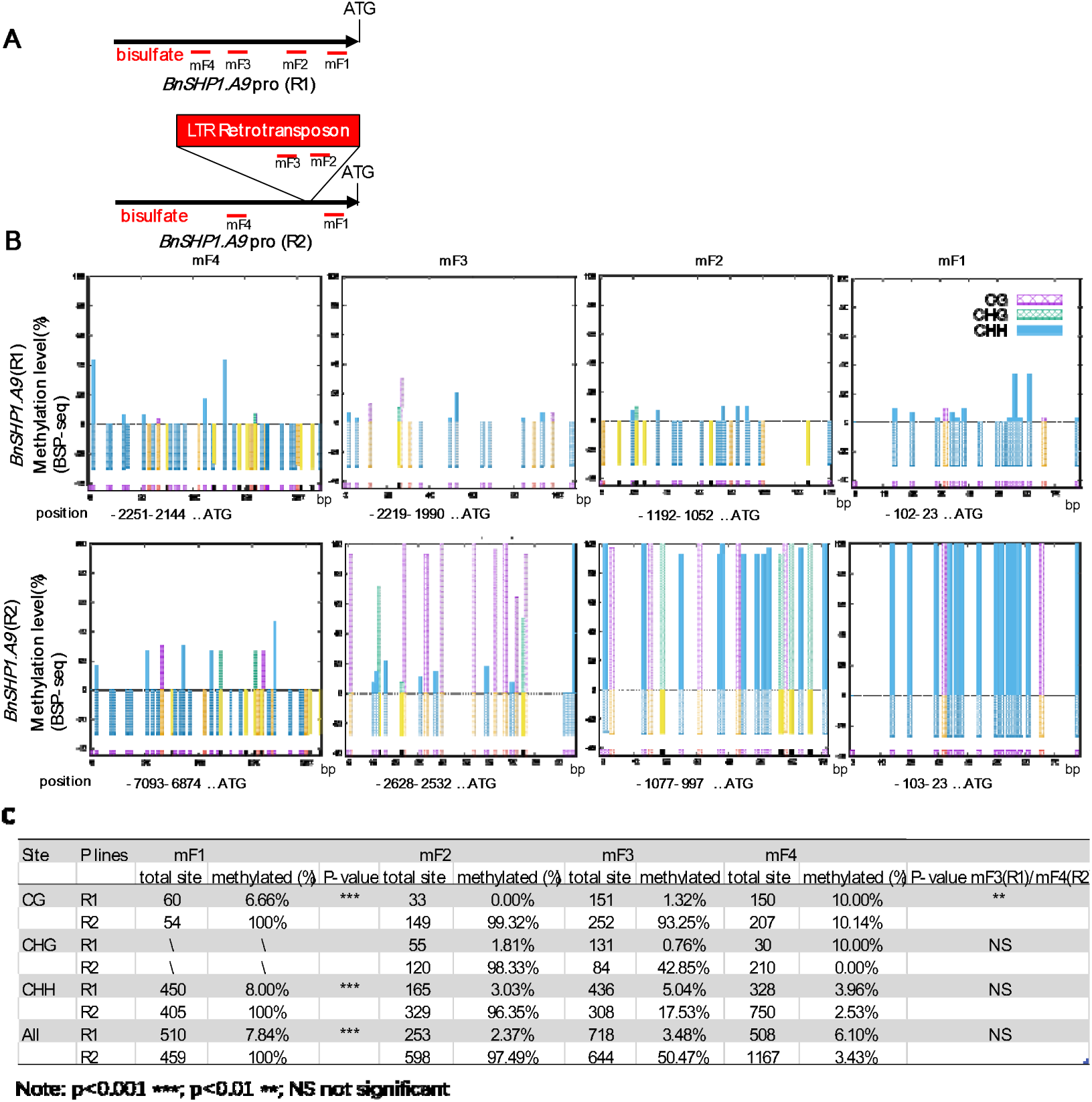
Methylation analysis of upstream promoter region of the *BnSHP1.A9* comparison between R1 and R2. **A** The positions of BSP analysis in the upper 5’ UTR region of *BnSHP1.A9* (black line) and 4803 bp *copia* LTR-retrotransposon insertion (red box) are illustrated. Red lines indicate the position of the bisulphite sequencing (BSP) analyses. **B** Methylation of cytosine residues in CG, CHG and CHH sites (purple, green and blue lines, respectively) was revealed by bisulphite sequencing of the four BSP regions. **C** Methylation data of cytosine residues in CG, CHG and CHH sites from the four BSP regions

### The LTR insertion correlates with pod shatter resistance in diverse oilseed rape germplasm

To verify the linkage between this LTR retrotransposon insertion and pod shatter resistance, we tested 135 diverse accessions with an allele specific diagnostic marker for LTR detection (BnSHP1.A9p1/p2, Table 1). The homozygous *BnSHP1.A9* (R1) allele (R1 specific, LTR^-^) and homozygous *BnSHP1.A9* (R2) allele (R2 specific, LTR^+^) were detected in 63.7% (86) and 15.6% (21) accessions, respectively, and heterozygous *BnSHP1.A9* (H) was found in 11.9% (16) accessions (Supplemental table 1). A significant association between the *BnSHP1.A9* (R2, LTR^+^) promoter allele and pod SRI was observed among the tested oilseed rape accessions (Figure 7). The average pod SRI of the lines with *BnSHP1.A9* (R2, LTR^+^) genotype was significantly higher than that of lines with *BnSHP1.A9* (R1, LTR^-^), indicating that *BnSHP1.A9* (R2, LTR^+^) allele could increase PSRI compared to *BnSHP1.A9* (R1, LTR^-^) allele.

**Figure 7.**
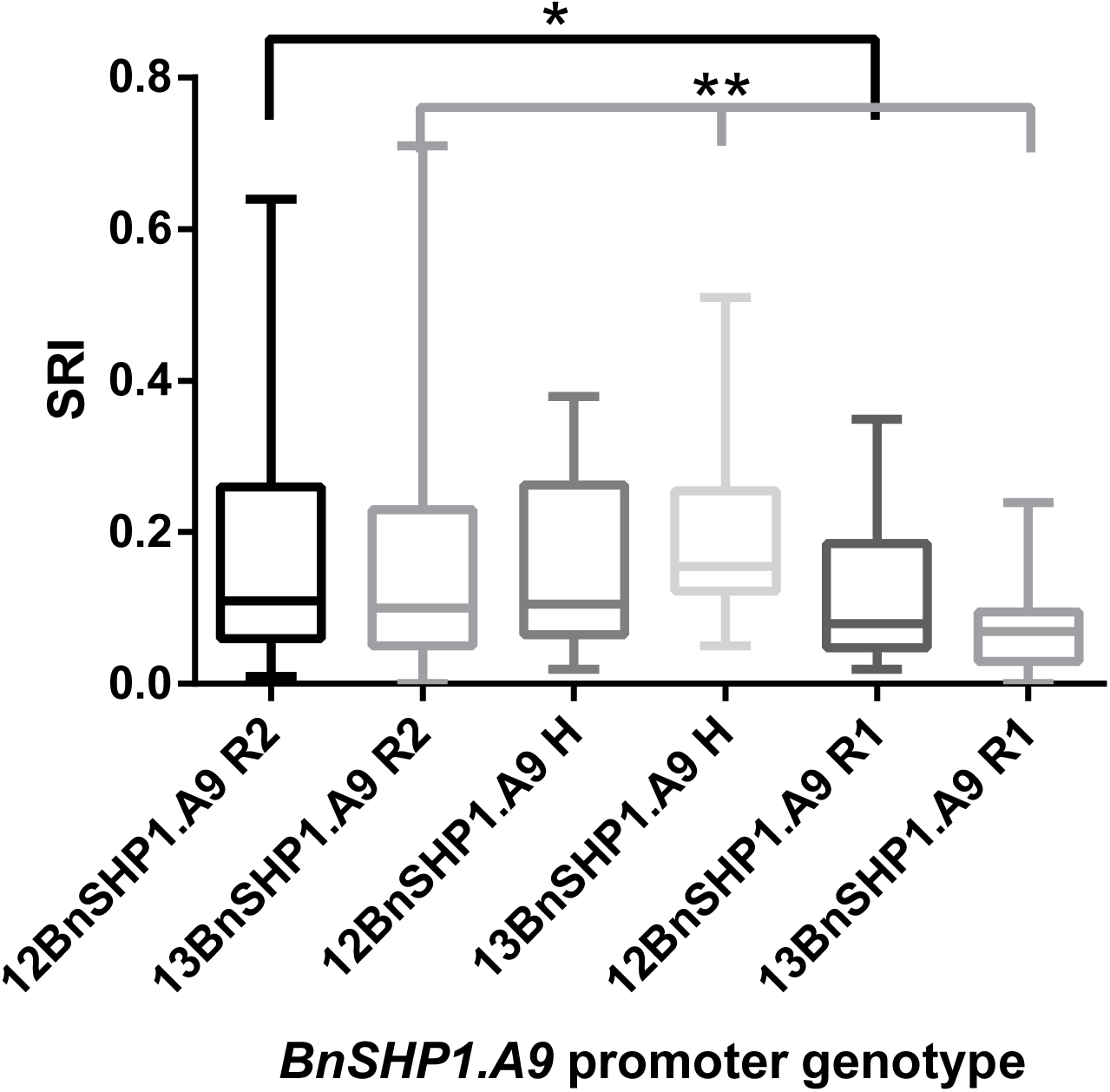
The distribution of *BnSHP1.A9* promoter allele in natural population and corresponding PSRI in 2012 and 2013 environments.

## Discussion

### *SHATTERPROOF 1* gene, *BnSHP1.A9* underlies pod shatter resistance

In this study, we delineated two QTL; *qSRI.A9.1* and *qSRI.A9.2* with large allelic effects, on chromosome A9 in a DH population from R1/R2. QTL accounting for large phenotypic variance of pod shatter resistance have been identified in various *B. napus* populations (Hu et al., 2012; Raman et al., 2014; Liu et al., 2016). Since a *SHP1* paralog of *A. thaliana* is located in the vicinity of the significant SNP markers associated with pod shatter resistance on chromosome A9 (Raman et al., 2014; Liu et al., 2016, this study), we were interested in determining whether *SHP1* indeed contributes to the genetic variation for pod shatter resistance in *B. napus* germplasm. Through genetic analyses (using a DH population from R1/R2 and a set of 135 diverse lines of oilseed rape breeding germplasm) and functional characterization via comparative expression analysis and transgenic approaches, we showed that the *SHP1* paralog, *BnSHP1.A9* underlies pod shatter resistance at *qSRI.A9.1* in oilseed rape.

In this study, we revealed that *BnSHP1.A9* is the functional gene regulating pod shatter resistance in R1/R2 DH population and in diverse *B. napus* lines. By overexpressing *BnSHP1.A9* cDNA from both R1 and R2 alleles, an average of 50% of PSRI decrease was found in T1 lines, thus confirming the function of *BnSHP1.A9* in pod shatter regulation. This functional analysis result is in accordance with the expression pattern of *BnSHP1.A9* in the R1 and R2 parental lines. In R2 which contributed positive effect to PSRI on this locus, the expression of *BnSHP1.A9* was repressed, indicating that the down-regulation of the target gene enhances PSRI. Overexpression of *BnSHP1.A9* only partly decreased the pod shatter resistance of R1. This can be attributed to other loci known to contribute positive effect in R1 for pod shatter resistance, such as the *qSRI.A9.2* as well as other *SHP1* or *SHP2* homologues which are known to act redundantly and control dehiscence zone differentiation (Liljegren et al., 2000). Although the expression difference of *BnSHP1.A9* exists in lines with contrast genetic effects of this locus, the allelic variation of *BnSHP1.A9 per se* is not the causal factor for the phenotypic variation of pod shatter resistance, as overexpression of both alleles of the *BnSHP1.A9* gene facilitated pod shattering in *B. napus*.

### LTR retrotransposon insertion in the upstream region regulates *BnSHP1.A9* epigenetically

Comparative and association analyses revealed that the LTR retrotransposon insertion was significantly associated with pod shatter resistance among 135 collected accessions, and in R1/R2 DH population. The *BnSHP1.A9* (R2 allele) promoter region, including LTR retrotransposon insertion was found to be highly methylated, which is responsible for the depression of *BnSHP1.A9* expression and the positive effect to pod shatter resistance phenotype. Transposable elements (TEs) are well known to play positive roles in generating genomic novelty and diversity in plants (Song et al., 2017). TEs are frequent found in *B. napus* genomes (Chalhoub et al., 2014; Sun et al., 2017) and have been implicated in DNA methylation and H3K9me2 modification (Eichten et al., 2012; Gent et al., 2013), altering gene expression both genetically and epigenetically (Cui and Cao, 2014). In fact, several oilseed rape genes related to morphological or physiological traits have evolved from TE insertions (Hou et al., 2012; Zhang et al., 2015; Gao et al., 2016; Shi et al., 2019). The current study implies that DNA methylation of the Copia-LTR insertion spreads to *BnSHP1.A9* cis-regulatory region. This epigenetic modification may change the accessibility of RNA polymerase II and transcription factors to the *BnSHP1.A9* promoter, and ultimately altering transcription patterns (Zhao et al., 2013).

*B. napus* is an allotetraploid originated from natural hybridization of *B. rapa* and *Brassica oleracea*. The LTR insertion in *BnSHP1.A9* promoter region showed 100% sequence identities with *B. rapa*. This suggests that the insertion of LTR retrotransposon event took place before the generation of *B. napus* as a species. However, only 15.6% of the natural *B. napus* population was found to contain the LTR insertion, which indicates that the LTR insertion might be lost in the process of domestication or breeding of rapeseed. As pod shattering is beneficial to seed release, the loss of LTR insertion is evolutionarily advantageous. This is further consistent with the theory that pod shatter resistance was selectively against during natural evolution or domestication.

We thus propose a simple model to explain the *BnSHP1.A9* dependent pod shatter resistance in *B. napus*. The non-LTR *BnSHP1.A9* (in R1, LTR^-^) is in a transcriptionally active state. DNA methylation of the LTR insertion in *BnSHP1.A9* promoter (in R2, LTR^+^) spreads to the transcription initiation region, thus converting *BnSHP1.A9* from an active state to a silenced state. In this study, hypermethylation of the *BnSHP1.A9* promoter region, mainly through CG and CHH methylations, appears to be the major epigenetic factor in the regulation of gene expression. It seems reasonable for plants to evolve such an epigenetic regulatory mechanism to gain functional variation for pod shatter resistance.

### Application of *BnSHP1.A9* gene for oilseed rape breeding

In this study we investigated the molecular basis of pod shatter resistance in oilseed rape utilizing natural variation in *B. napus* germplasm and showed that a single gene, *BnSHP1.A9,* controls genetic variation for pod shatter resistant at *qSRI.A9.1* locus. Our research provides two gene-specific markers, one co-dominant marker (BnSHP1.A9IF4) detecting sequence difference within the CDS of *BnSHP1.A9*, and the other co-dominant marker (BnSHP1.A9P1/P2) detecting presence/absence variation (PAV) of LTR insertion in *BnSHP1.A9* promoter region. Both markers can be applied for the efficient selection of this QTL for pod shatter resistance in *B. napus* breeding programs starting from early generations. Previously, there were only linked markers available for marker assisted selection for pod shatter resistance in *B. napus* to PSRI QTL reported and could be used for breeding (Raman et al., 2014; Liu et al., 2016). Our two gene-specific markers developed in this study could be used to further improve the resistance potential by introducing the *BnSHP1.A9* (R2) allele with positive effect into lines containing the *BnSHP1.A9* (R1) allele for genetic improvement. These markers could be easily assayed via conventional agarose gel electrophoresis and high throughput capillary electrophoresis platforms. The sources of pod shatter resistance identified herein can be used for introgression of favorable alleles and enrichment of alleles in the breeding germplasm.

Recently CRISPR-Cas9 genome editing and other genetic transformation platforms have become available for oilseed rape improvement (Zaman et al., 2019). Editing IND and ALC genes has improved the pod shatter resistance in *B. napus* (Braatz et al., 2018a; b). Our results clearly revealed that down-regulation of *BnSHP1.A9* could increase PSRI. Genome editing can not only mutate *BnSHP1.A9*, but also other functionally redundant homologous of *SHP1*, as well as the homologous of *SHP2*. Knockdown the multiple copies of functional redundant genes or homologues has more effect on phenotypic variation. However, these approaches are difficult to deploy commercially due to the legal restrictions on GMO crops in some countries, such as Europe. In countries that are open to gene edited crops, our finding on the down-regulation of the *BnSHP1.A9* having a positive effect on pod shatter resistance can be used to manipulate the gene expression by mutation of this gene and its homologs for the improvement of pod shatter resistance in *B. napus*.

In conclusion, we showed that a *SHP1* paralog, controls pod shatter resistance at qSRI.A9.1 QTL in the DH population from R1/R2 and in a diversity panel of oilseed rape. Our results suggest that upstream promoter region of *BnSHP1.A9* controls pod shatter resistance, rather than the coding sequence. This study provides a novel source of germplasm, gene-specific markers and insights on molecular basis of *SHP1* mediated resistance to pod shatter in oilseed rape. These resources will facilitate the genetic improvement of pod shattering resistance in oilseed rape.

## DECLARATIONS

### Ethics approval and consent to participate

Not applicable

### Consent for publication

Not applicable

### Authors’ contributions

QH and HR designed the research. JL, RZ, WH, WW, DM and TC performed genetic analysis and field research. RZ performed genetic transformation and DNA methylation analysis, JL and RZ analyzed data. YQ and RR performed SHP1 analysis using capillary electrophoresis, JL wrote the first version of the manuscript. All authors reviewed the manuscript.

