## Supplemental Figure for "An Epigenetic LTR-retrotransposon insertion in the upstream region of *BnSHP1.A9* controls quantitative pod shattering resistance in *Brassica napus*"

Supplemental figure 1. *BnSHP1.A9* expression analysis of leaves in the 5 T1 lines by RT-PCR. *BnActin* gene was used as an internal control for relative expression analysis. 1, 2, 3: three individual plants of T18: 4, 5, 6: T24; 7, 8, 9: T26; 10, 11, 12: T33; 13, 14, 15: T9.

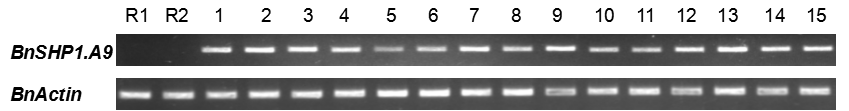
